# When are bacteria really gazelles? Comparing patchy ecologies with dimensionless numbers

**DOI:** 10.1101/2021.10.15.464607

**Authors:** Samuel S. Urmy, Alli N. Cramer, Tanya L. Rogers, Jenna Sullivan-Stack, Marian Schmidt, Simon D. Stewart, Celia C. Symons

**Affiliations:** Monterey Bay Aquarium Research Institute, 7700 Sandholdt Rd., Moss Landing, CA 95039, USA; School of the Environment, Washington State University, Pullman, WA 99163, USA; NOAA Southwest Fisheries Science Center, 110 McAllister Way, Santa Cruz, CA 95060, USA; Department of Integrative Biology, Oregon State University, 3029 Cordley Hall, Corvallis, OR 97331, USA; Department of Integrative Biology, University of Texas at Austin, Texas 78712, USA; Cawthron Institute, 98 Halifax St East, Nelson 7010, New Zealand; University of California, Irvine, Department of Ecology and Evolutionary Biology, 469 Steinhaus Hall, Irvine, CA 92697, USA

**Keywords:** Comparative ecology, Dimensional analysis, Interactions, Model systems, Patchiness, Scale

## Abstract

From micro to planetary scales, spatial heterogeneity—patchiness—is ubiquitous in ecological systems, defining the environments in which organisms move and interact. While this fact has been recognized for decades, most large-scale ecosystem models still use spatially averaged “mean fields” to represent natural populations, while fine-scale, spatially explicit models are mostly restricted to particular organisms or systems. In a conceptual paper, Grünbaum (2012, *Interface Focus* 2: 150-155) introduced a heuristic framework, based on three dimensionless ratios quantifying movement, reproduction, and resource consumption, to characterize patchy ecological interactions and identify when mean-field assumptions are justifiable. In this paper, we calculated Grünbaum’s dimensionless numbers for 33 real interactions between consumers and their resource patches in terrestrial, aquatic, and aerial environments. Consumers ranged in size from bacteria to blue whales, and patches lasted from minutes to millennia, spanning spatial scales of mm to hundreds of km. We found that none of the interactions could be accurately represented by a purely mean-field model, though 26 of them (79%) could be partially simplified by averaging out movement, reproductive, or consumption dynamics. Clustering consumer-resource pairs by their non-dimensional ratios revealed several unexpected dynamic similarities between disparate interactions. For example, bacterial *Pseudoalteromonas* exploit nutrient plumes in a similar manner to Mongolian gazelles grazing on ephemeral patches of steppe vegetation. Our findings suggest that dimensional analysis is a valuable tool for characterizing ecological patchiness, and can link the dynamics of widely different systems into a single quantitative framework.

## Introduction

Most ecosystems are patchy across a range of spatial and temporal scales, and, consequently, most ecological interactions occur in patchy environments (Hutchinson 1961; Wiens 1976; Levin 1994; Lawton 1999). In the ocean, predators search for shifting schools of fish (Weimerskirch *et al*. 2005; Benoit-Bird *et al*. 2013). In the forest, grazers move from plant to plant or stand to stand (de Knegt *et al*. 2007). And at the micro-scale, bacteria search for organic particles and plumes of dissolved nutrients (Stocker 2012; Hellweger 2018; Yawata *et al*. 2020). The uneven distribution of resources means that consumers (i.e., predators, herbivores, or detritivores) often experience resources far from their mean density. Patchiness and heterogeneous dynamics have been recognized as critical issues in ecology for decades (Durrett & Levin 1994; Codling & Dumbrell 2012) and form the basis of many ecological theories (Chesson 2000; Davis *et al*. 2005). However, ecologists still struggle to incorporate them into our mental, mathematical, and management models.

Two main challenges have prevented the full integration of patchiness into ecology and management. First, patches are by definition localized in space and time, so incorporating them requires temporally and spatially explicit models. Spatially explicit models are more theoretically complex, computationally expensive, and difficult to use than non-spatial models (Morozov & Poggiale 2012; DeAngelis & Yurek 2017). Consequently, most researchers still rely on “mean-field” models, assuming that spatially variable quantities can be represented by a single average value (Codling & Dumbrell 2012). Despite their simplicity and wide use, mean-field models are often inadequate descriptions of real ecological dynamics (Priyadarshi *et al*. 2019). They are sensitive to functional responses at low population densities, often require arbitrary assumptions of ecological thresholds, and can erroneously predict competitive exclusions or extinctions in environments where real populations persist (Hutchinson 1961; Murray 1989; Grünbaum 2012). While mean-field approximations may be appropriate in some situations, they must be justified based on specific hypotheses and scales of ecological processes.

The second challenge arises because ecological patches occur across a vast range of physical scales. Patchiness is present in almost all systems at *some* scale, and many statistics and metrics exist to quantify it (Fortin 1999). However, it is less clear how to determine which patches, at which scales, are relevant for which organisms. Moreover, when a patchy process or interaction has been described for one species or system, it is often unclear which, if any, of its conclusions apply more generally. Such studies can remain isolated in their (sub)discipline’s literature as apparent special cases, with no way to identify similar dynamics in other taxa or environments.

### Box 1

Dimensional analysis and dynamic similarity

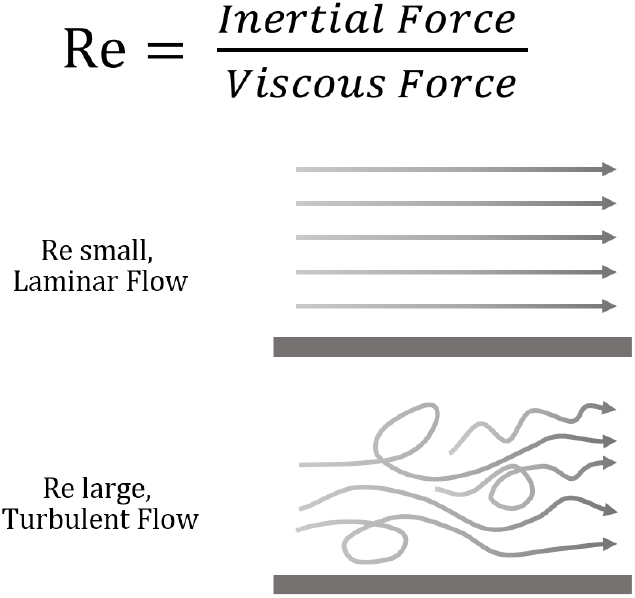

Dimensional analysis is a technique for simplifying mathematical representations of physical problems. The key to this method is to reduce the number of parameters required to describe a system by combining the original variables into a smaller number of dimensionless ratios—i.e., ratios where all the physical units cancel out, leaving only a pure number expressing the relative importance of its components. Despite widespread use in the physical sciences, dimensional analysis is still relatively rare in ecology (a notable exception being the relationship between growth, *K*, and mortality *M*, (Beverton & Holt 1959; Charnov 1990; Charnov *et al*. 1991)).

Dimensional analysis can provide *quantitative* insight by identifying the importance of different variables. It can also provide *qualitative* insights by identifying fundamental properties of systems with similar dynamics. A well-known physical example is the Reynolds number (Re), which gives the ratio of inertial to viscous forces within a fluid. Quantitatively, if Re >> 1 a fluid dynamicist can safely drop all frictional terms from their equations, even though some friction is, of course, always present. Qualitatively, flows with Re greater than about 5 × 10^5^ are likely to be turbulent, and thus fundamentally different from the laminar flows prevalent at lower Re. Since Re is unitless, this pattern holds true regardless of the particular fluid and physical scale. Thus, Re is as useful for describing the biomechanics of plankton as it is for designing airplanes.

Systems are *dynamically similar* when the relative importance of their attributes are the same – i.e., the ratios between variables are the same. Dynamically similar systems can serve as models for one another, even if their absolute scales differ widely.

One approach capable of addressing both of these challenges is dimensional analysis, as it facilitates comparisons between processes, regardless of scale (Stephens & Dunbar 1993; Horne & Schneider 1994, Box 1). In a theoretical paper, Grünbaum (2012) proposed a dimensional analysis of interactions between consumers and their resource patches. In this framework, the resource landscape is simplified as a collection of discrete patches which are separated by a characteristic distance, persist for a characteristic time span, and contain a characteristic density of resources. Consumers are distinguished by their typical movement rates, generation times, and consumption rates (Figure 1). From these parameters, three dimensionless ratios can be calculated: the Frost, Strathmann, and Lessard numbers (Fr, Str, and Le, respectively; Table 1). These ratios compare the duration of an undisturbed patch to the typical time required for consumers to locate a patch (Fr), to reproduce (Str), and to consume a patch (Le). The three ratios summarize the importance of movement, reproduction, and resource depletion to the consumer-resource interaction, independent of the absolute scales and rates involved.

**Table 1.** Dimensionless ratios from Grünbaum (2012). In each ratio, *T* represents the typical patch duration, *T*_*search*_ is the time required for a consumer to locate a patch, *T*_*repro*._ is the consumer’s generation time, and *T*_*cons*._ is the time required for consumers to deplete a patch. All of these durations are “characteristic time scales,” accurate to an order of magnitude but not necessarily more so.

| Dimensionless ratio | Description | Ratio $\gg 1$ | Ratio $\ll 1$ |
| --- | --- | --- | --- |
| Frost number<br>$Fr = T / T_{search}$ | The importance of forager movement to resource patch occupancy | Consumers find patches easily and spend most time inside them | Consumers spend most time searching for patches |
| Strathmann number<br>$Str = T / T_{repro.}$ | The importance of forager reproduction to resource patch occupancy | Consumer populations can grow inside patches | Consumers must visit many patches each generation |
| Lessard number<br>$Le = T / T_{cons.}$ | The importance of forager consumption to resource patch persistence | Consumers actively deplete patch resources | Patch resource densities not affected by consumption |

**Figure 1.**
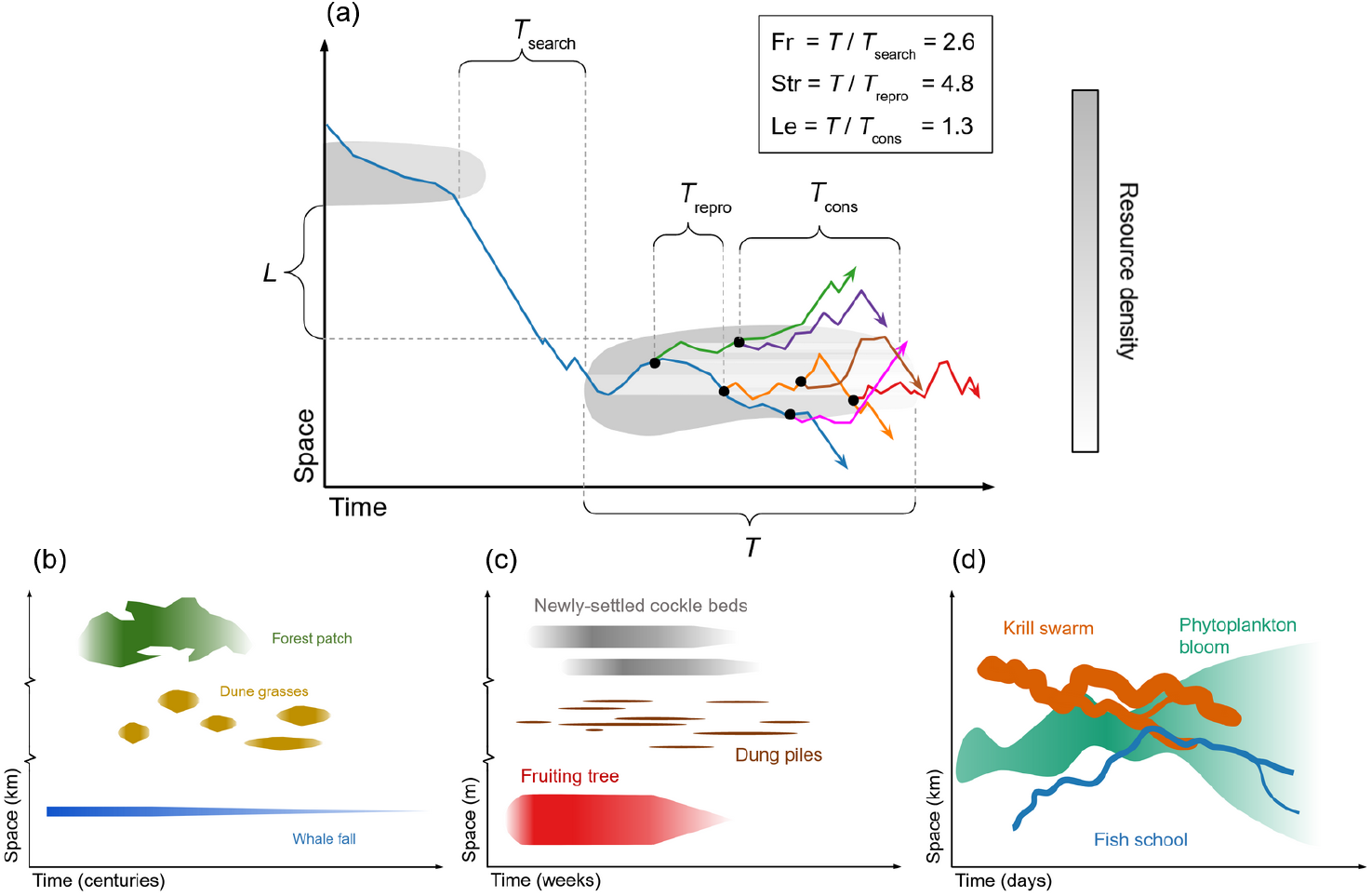
Conceptual space-time depictions of patchy consumer-resource interactions. (a) Hypothetical interaction between consumers (solid colored lines) feeding on resource patches (grey regions). As time (horizontal axis) progresses, consumers move on a simplified one-dimensional spatial domain (vertical axis), seeking resource patches. When they enter a patch, they begin to deplete it, indicated by lightened shading to the right of consumer trajectories (i.e., after their passage). Black dots represent reproduction events, and hence the birth of new consumers. Brackets show the characteristic duration of patches (*T*) and the distance between them (*L*), as well as the time scales for consumers to locate a patch (*T*_search_), reproduce (*T*_repro_), and consume a patch (*T*_cons_, assuming they are at a “typical” density within it). The box in the upper right gives the ratios of the interaction’s time scales as its dimensionless Frost (Fr), Strathmann (Str), and Lessard (Le) numbers. Panels (b), (c), and (d) depict the different physical scales of various resource patches. (b) Forest stands, dune grass patches, and whale falls persist for tens or hundreds of years and are spaced at kilometer scales. (c) Trees bear ripe fruit, dung piles remain favorable for dung beetles, and newly settled cockles are small enough for crabs to eat for weeks, with spatial scales of meters. (d) Phytoplankton blooms persist for days to weeks, while zooplankton and fish aggregations last from hours to days, with spatial scales of km.

The most practical application of these ratios is to determine which consumer-resource interactions can be approximated using a mean-field model. If Fr, Str, and Le are “small” (i.e., << 1), then consumer movement, reproduction, and consumption operate on time scales much longer than those of individual resource patches. Consumers encounter patches randomly and integrate over many patches in time, justifying the mean-field assumption. On the other hand, if any of the ratios are *not* “small,” then the interaction is “functionally patchy,” and must be treated as such. For instance, if Fr >>1, consumers will spend most of their time inside patches, experiencing local resource densities well above the average value. In circumstances like these, mean-field assumptions underestimate the resources available to the consumers (Lasker 1975; Benoit-Bird *et al*. 2013). Grünbaum (2012) also proposed several conjectures for patchy interactions. For instance, consumer populations should only persist if they can locate patches easily, reproduce within patches, or both. This leads to the prediction that all interactions should have at least one of Fr or Str > 1. Relatedly, if either movement or reproduction is difficult or costly for a consumer species, evolution should select for movement or reproductive rates that are “just fast enough” to raise Fr or Str above the critical value of ≈1.

Despite their conceptual power and potential utility, to our knowledge these dimensionless ratios have never been empirically calculated, which was the objective of this study. We conducted a literature review and meta-analysis of patchy consumer-resource interactions across a wide range of taxa, scales, and ecosystems. We obtained values for consumer rates and the spatial and temporal scales of their resource patches, and used these to calculate Frost, Strathmann, and Lessard numbers for each interaction. Using these ratios, we determined which interactions were functionally patchy and which could be approximated using mean-field assumptions, and we evaluated several of Grünbaum’s conjectures. Finally, we identified consumer-resource interactions that were dynamically similar — that is, close together in Fr-Str-Le space. This approach identified commonalities which might otherwise be obscured by differences in taxon, environment, or spatiotemporal scale, shifting the focus from individual species towards generalized interactions between consumers, resources, and their environments. The ultimate goal of such an analysis is to enable more parsimonious models and a deeper, more general understanding of patch dynamics in ecology.

## Methods

We took a meta-study approach to calculate values of Grünbaum’s dimensionless ratios for consumers feeding on patchy resources. For this analysis, a “patch” was defined as a spatially and temporally bounded region, within which a resource is dense enough for profitable exploitation by a consumer (Charnov 1976; Wiens 1976, Figure 2). Some researchers have used “patchiness” as a loose synonym for “heterogeneity,” including spatial gradients (e.g., (Arditi & Dacorogna 1988), but we required patches to contain a local maximum of the resource’s density. We restricted our analysis to trophic interactions, where exploitation of the resource involved energy transfer to the consumer. For example, a hydrothermal vent is a resource patch for chemosynthetic organisms, because they use the vent fluids as a source of chemical energy. In contrast, a lake is not a resource patch for a fish—rather, it is a habitat.

To calculate values for Fr, Str, and Le, we searched the literature for data on specific consumer-resource pairs. Pairs were selected to reflect a diversity of systems, based on author expertise and availability of necessary data. Literature searches were performed using Google scholar and Web of Science for the parameters of interest (Table 2). Search terms included “speed”, “turning interval”, “movement”, “growth rate”, “reproduction”, “consumption rate”, “foraging”, and “patch”. We took values directly from published sources when possible, though some values were estimated based on published ranges or the interpretation of published figures. In the rare cases where data were not available, parameters were estimated either via the authors’ expertise or consultation with experts. All raw values used for the analysis, with their sources, are tabulated in Appendix S1. While our selection of consumer-resource interactions cannot be considered random or unbiased, we made conscious efforts to represent marine, freshwater, and terrestrial environments; microbial, invertebrate, and vertebrate taxa employing carnivorous, herbivorous, and detrital feeding strategies; and patches which were stationary, passively moving, and actively moving.

**Table 2.**
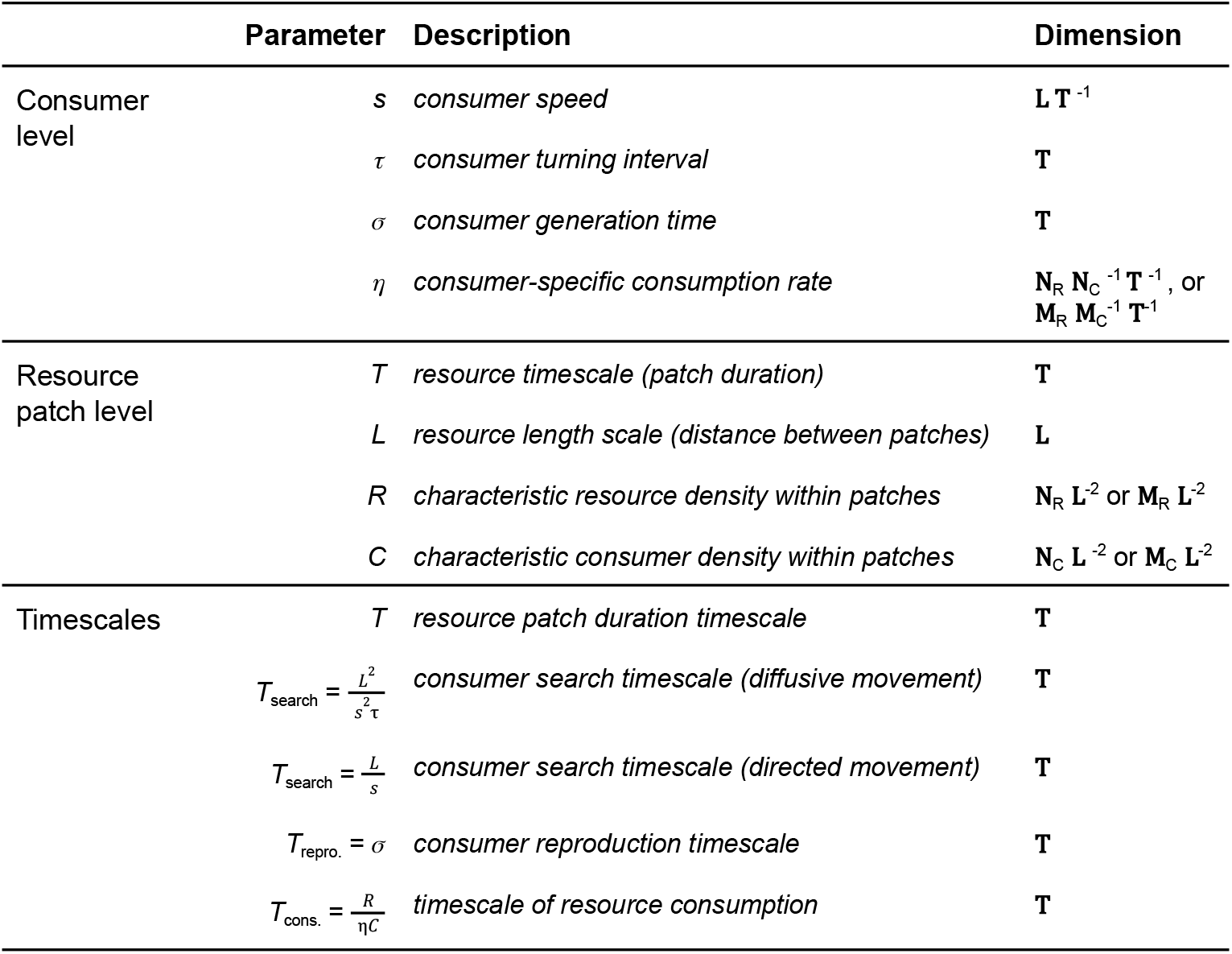
Parameters and rates obtained from the literature to develop data-derived Frost, Strathmann, and Lessard numbers. The physical dimensions of each quantity are denoted as **T** (time), **L** (length), **N** (number of individuals), and **M** (biomass), with the subscripts “R” and “C” indicating “resource” or “consumer.” (Adapted from Table 1 in Grünbaum 2012.)

These “characteristic” rates, scales, and densities for consumers and resource patches were necessarily broad-brush descriptions; selecting a single value based on the available literature was often challenging and somewhat arbitrary. Fortunately, for their intended use as descriptive heuristics, the dimensionless numbers did not require precision greater than 1-2 significant digits. With better data and domain knowledge it would be possible to calculate them more precisely, but was beyond the scope of this paper. In practice, the dimensionless ratios varied across many orders of magnitude (see Results and Discussion), so greater precision would not have qualitatively changed the results.

To allow for comparisons, we converted all rates and quantities to base SI units (seconds, meters, and grams) and checked for dimensional consistency. Values for Fr, Str, and Le were then calculated for each consumer-resource pair. We calculated two Frost numbers for each consumer: one based on an assumption of diffusive (i.e., random walk) movement, and one on directed-line (i.e., ballistic) movement. These two movement types are unrealistically simple, but define reasonable bounds for the range of more complex movement types used by real organisms.

To determine when systems are functionally patchy and to visualize the potential impact of that patchiness on consumer-resource dynamics, we developed four-quadrant diagrams similar to those in Grünbaum (2012). These diagrams provide a graphical assessment of patch dynamics and the importance of consumer movement (Fr), reproduction (Str), and patch consumption rate (Le) to trophic interactions. These three ratios can be visualized as a point in three dimensions; for readability, we display the ratios on logarithmic axes, and examine this 3-D point cloud projected on only two axes at a time. These figures also let us compare our results with predictions from Grünbaum (2012). We also calculated the Pearson product-moment correlation between each pair of dimensionless numbers.

To determine which consumer-resource pairs were dynamically similar, we performed a cluster analysis based on the Euclidean distance between the log_10_ Fr, Str, and Le ratios for each consumer-resource pair with complete agglomeration. The dendrogram was partitioned into five discrete clusters using a k-medioids algorithm (Reynolds *et al*. 2006; Schubert & Rousseeuw 2019) within the ComplexHeatmap R package (Gu *et al*. 2016). This number of clusters was chosen based on the “elbow method” (Thorndike 1953, Figure S2) and our subjective assessment of the level of clustering that generated biologically meaningful groups. Within each cluster, we examined each interaction’s ecosystem, consumer type, patch mobility, and consumer/resource size ratio, to explore whether these characteristics were associated with particular types of consumer-resource dynamics. All data and associated code for this project are available at https://github.com/allicramer/EcologicalPatchiness.

## Results and Discussion

We analyzed a total of 33 interactions between consumers and their resource patches. Of these, 12 took place in terrestrial environments, 18 in marine environments, and 3 in freshwater (Table S1). The consumers ranged in average body size from 0.5 μm to 22 m, and in mass from 10 picograms to 5 tons. They included representatives of multiple trophic strategies (11 herbivores, 17 carnivores, and 5 detritivores) and body plans (2 microbes, 13 invertebrates, and 18 vertebrates). In most cases resource patches were composed of more than one organism; these covered a similarly large range of body sizes (Figure 3a). The resource patches lasted anywhere from 5 minutes (groups of seals swimming from their haul-out to deep water) to 1500 years (whale falls being buried by sediment), and were separated by distances ranging from 4 cm (nutrient patches from lysing phytoplankton) to 55 km (schools of zooplankton and fish in the open ocean) (Figure 3b). When the consumers’ search, reproduction, and patch-depletion times were normalized by the durations of their resource patches, the resulting Frost, Strathmann, and Lessard numbers ranged across more than 10 orders of magnitude (Table 3, Figure 4). Despite the huge diversity of biology and range of scales considered, the dimensional analysis framework allowed us to identify a number of generalities, trends, and similarities across all the interactions.

**Table 3.**
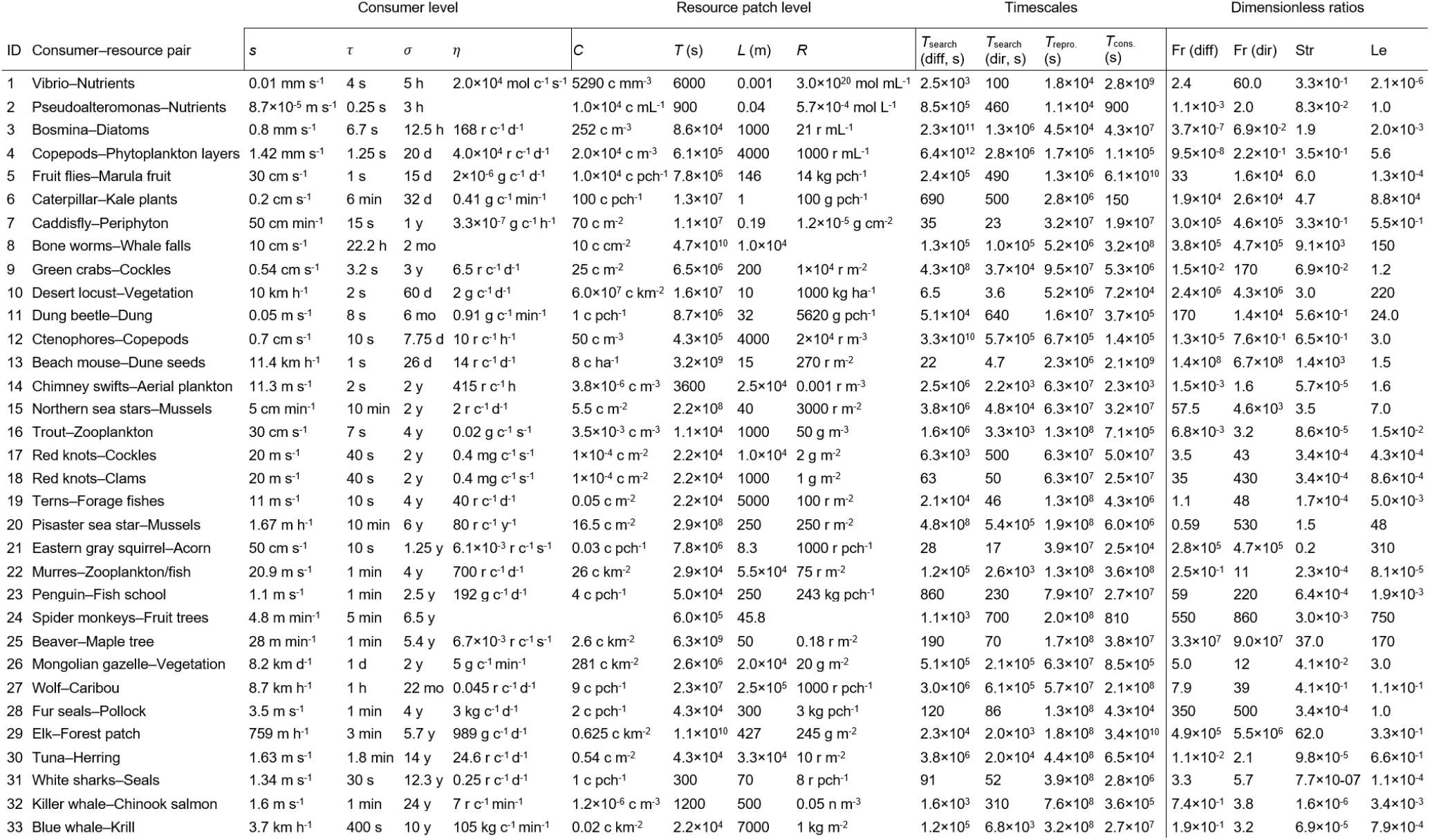
Rates and dimensionless ratios for all consumer-resource pairs. Parameters are defined in Table 2; see Table S1 for a more detailed version of this table with references. Consumer and resource quantities are presented in their original units from the primary literature, which vary from interaction to interaction. For instance, the “currency” of an interaction could be expressed in individuals or biomass, densities could be areal or volumetric, etc. However, when plugged into the equations in Table 2, the quantities within each row are dimensionally consistent. Abbreviations for units: m = meters, g = grams, s = seconds, min = minutes, h = hours, d = days, mo = months, y = years, ha = hectares, c = consumer individuals, r = resource individuals, mol = molecules, pch = patch, diff = diffusive, dir = directed.

**Figure 3.**
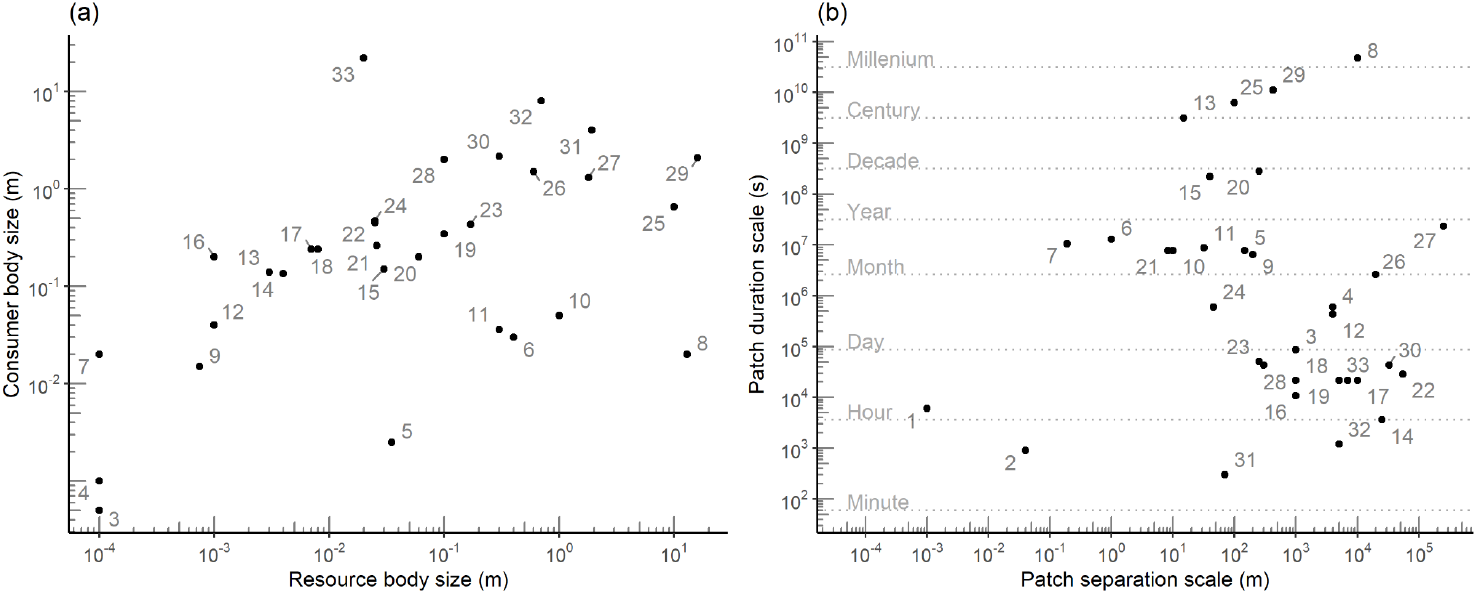
The sizes of consumer and resource organisms, and the spatio-temporal scales of resource patches, spanned many orders of magnitude. (a) Body size of consumer organisms (y-axis) plotted against the size of an individual resource item. Numbered labels refer to the interaction IDs in Table 3, each containing a consumer organism and a corresponding resource (Note that the two bacterial interactions are excluded, since “organism size” is poorly defined for a dissolved nutrient resource). (b) Resource patches spanned approximately eight orders of magnitude in their spatial and temporal scales. Each point represents the resource patch within an interaction, labeled as in (a). The horizontal axis represents the typical distance scale separating patches in space, ranging from ∼1 mm (microscale nutrient plumes, interaction 1) to ∼250 km (caribou herds, interaction 27). The vertical axis represents the patches’ typical duration, ranging from ∼5 min (seal groups, interaction 31) to ∼1,500 years (whale falls, 8).

**Figure 4.**
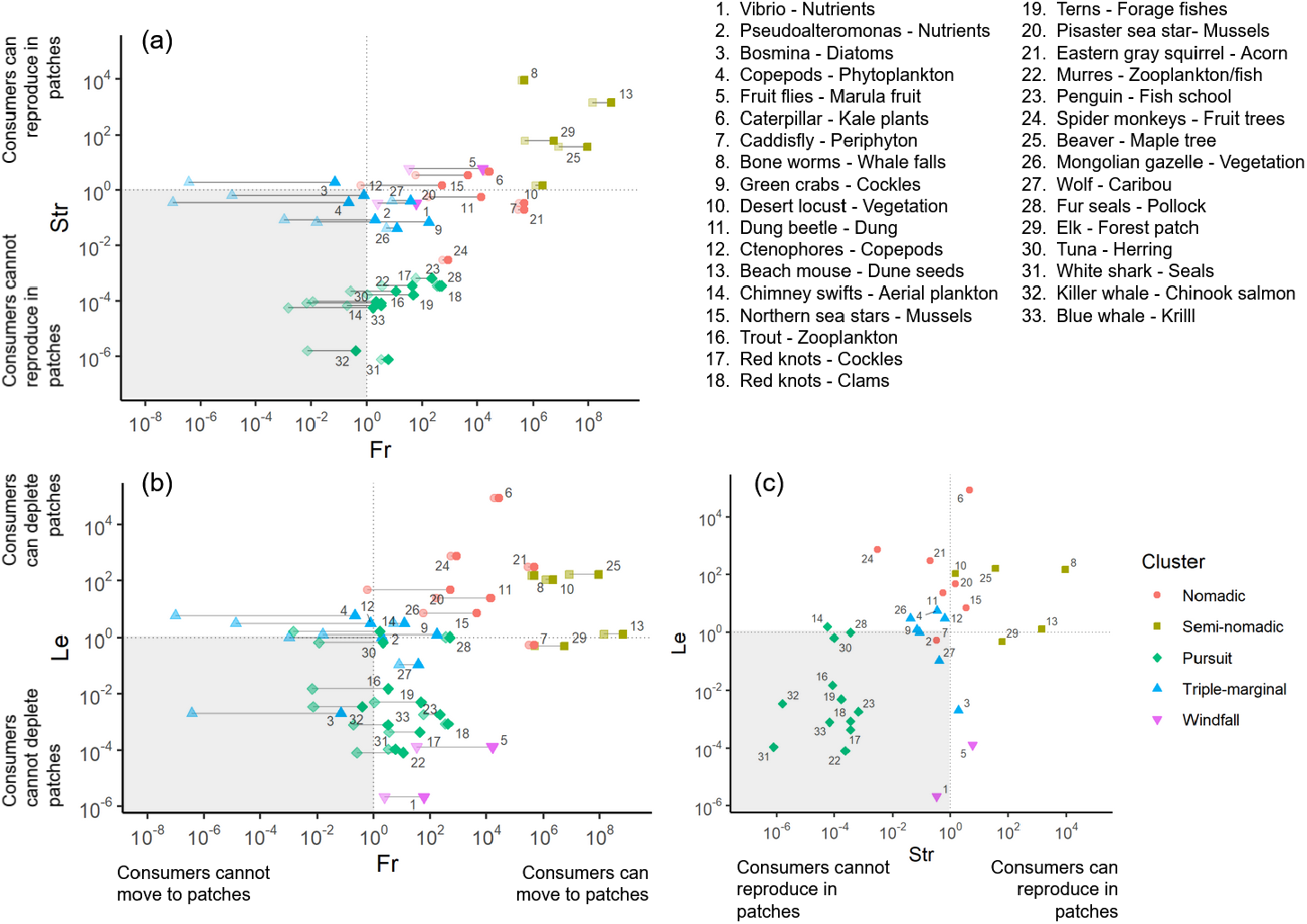
All consumer-resource interactions plotted on logarithmic axes in dimensionless Frost-Strathmann (a), Frost-Lessard (b), or Strathmann-Lessard (c) space, showing the relative importance of movement (Fr), reproduction (Str), and depletion (Le) in driving patch dynamics.. In (a) and (b), two points, connected by a line, are plotted for each consumer-resource pair: the lighter point is the Frost number assuming diffusive (i.e. random walk) movement, while the filled point is the Frost number for directed movement. The horizontal and vertical grey lines show the critical values, where Fr, Str, and Le equal 1. The shaded regions on each plot indicate interactions which are not functionally patchy with respect to each pair of dimensionless ratios. For interactions in these regions, the mean-field simplification may be (partially) justifiable. Point shapes and colors represent clusters of similar interactions identified via a k-medioids analysis; see Figure 5 and text for details.

### All interactions were functionally patchy

Every consumer-resource interaction considered here was functionally patchy with respect to movement, population growth, or consumption. All interactions had at least one of the three dimensionless ratios greater than 1, and in 25 out of the 33 pairs, at least one of the ratios was > 10 (Table 3). Seven of the 33 consumer-resource interactions (21%) had all three numbers > 1, indicating that patch dynamics had a significant influence on all of these aspects of consumer-resource ecology. Given this, a model using a strict mean-field assumption would not be justifiable for any of these interactions. While our deliberate selection of systems with patchy resources precludes generalizing this result to all consumer-resource interactions, it still stands as a warning to ecologists dealing with foraging movements, consumption/predation, or trophic transfer rates: if a system looks like it *might* have patchy dynamics, then they probably shouldn’t be ignored.

Even though patch dynamics could not be ignored entirely in any of the consumer-resource pairs, the majority (26 out of 33 interactions, or 79%) had at least one dimensionless ratio < 1, indicating that at least one of the processes (movement, reproduction, or consumption) *could* be simplified using a mean-field approximation. As a concrete example, consider interaction 19, between common terns *Sterna hirundo* and forage fishes. Terns prey on schools of small fishes forced to the surface by underwater predators (Safina & Burger 1985) or physical flows (Urmy & Warren 2018). Surface schools are typically separated by several km and remain available for up to a few hours. This interaction is characterized by a Fr_dir_ of 48, indicating that the terns’ rapid flight allows them to spend more time over fish patches than over empty water. When modeling trophic transfer between fishes and terns, estimates of energy flow based on the mean prey density (e.g., measured by a fishery survey) would be biased low. A more reasonable estimate would be parameterized based on the mean prey density *within* schools, perhaps with second-order effects to account for search times and other foraging dynamics. At the same time, terns visit many patches per generation (Str = 1.7 × 10^−4^), so if a parameterized energy transfer function were developed, averaging it over days, weeks, or even an entire breeding season *would* be justifiable. Finally, the low Lessard number (Le = 0.005) indicates that terns rarely deplete patches, so models of this interaction can treat the density and distribution of their prey as endogenous variables (cf. Urmy 2021).

Analogous simplifications can be identified for most of the consumer-resource interactions (Table 4). In general, low values of the Frost number indicate that spatial mean-field models may be appropriate, while high values mean they are not. As always, spatial mean-field models are highly dependent on scale; interactions with a high Frost number may be modeled based on mean densities *inside* patches. Similarly, low values of the Strathmann number suggest temporal averaging may be acceptable. With Str >> 1, many generations of consumers can reproduce in each patch and the appropriateness of mean-field models depends on the impacts of those generations (i.e., Le). If the consumers do not deplete the patches (i.e., Le << 1), the interaction is essentially a traditional metapopulation (Hanski 1998). On the other hand, if Le > 1, there will be a more complex balance between patch appearance, colonization, and depletion, similar to the susceptible-infected-recovered dynamics familiar from epidemiological models (Kermack *et al*. 1927).

**Table 4.** Guidance for modelling approaches based on dimensionless ratios.

| | Ratio $\ll 1$ | Ratio $\sim 1$ | Ratio $\gg 1$ |
| --- | --- | --- | --- |
| Fr | Consumers spend most time outside patches. Encounter and trophic transfer rates ∝ average densities of consumer and resource *over entire region*. Mean-field assumption may be ok. | Movement and foraging behaviors must be modeled or parameterized explicitly. | Consumers spend most of their time inside patches. Encounter and trophic transfer rates ∝ average densities of consumer and resource *within patches* (by definition, higher than spatial average). |
| Str | Consumers visit many patches per generation. Temporal averaging ok. | Multi-trophic population dynamics à la 2-species Lotka-Volterra, must be modeled or parameterized explicitly. | Multiple generations reproduce within each patch. If $Le \ll 1$, approaches metapopulation dynamics. |
| Le | Consumers have negligible effect on duration or density of resource patches, which appear and disappear independently of consumers. Feedbacks between consumer and resource density can be ignored. | Consumers may deplete resource patches. Effects of movement, reproduction, dispersal, and/or death must be modeled or parameterized to account for persistence of populations. If $Le \gg 1$, resource patches deplete quickly. | |

If any of the ratios are ∼1, simple approximations overlook crucial aspects of the system. In these cases, the relevant dynamic process(es) must be modeled or parameterized explicitly. Dimensionless ratios such as Fr, Str, and Le provide a simple way to check whether complex models are required. Crucially, they also provide a means to develop those models in a more general way. Models expressed in terms of dimensionless ratios benefit from a reduced parameter space and often express a problem in its simplest possible form (Stephens and Dunbar 1993). Additionally, in such models the specific units attached to the variables are irrelevant and the essential dynamics of the system become clear. Connections between dynamically similar systems can be identified independent of their absolute size or scale.

### Assessing Grünbaum’s conjectures for organisms in patchy environments

Across all interactions, the directed Frost and Strathmann numbers were positively correlated (r = 0.68, p < 0.001). Most interactions fell in the upper-right three quadrants in Fr-Str space (Figure 4a), indicating that consumers would be able to find and exploit patches through movement, reproduction, or both. For most consumers, the Frost number increased dramatically when directed rather than diffusive movement was assumed. Nine consumers would *only* be able to occupy patches if they used directed movement: if they relied on random search behavior, their theoretical search time would be longer than a patch’s typical duration. Figure S1 gives an alternate visualization of critical Frost numbers following the scheme of Grünbaum’s (2012) Figure 1a; this presentation suggests all consumers would be able to access their patches. Two consumer-resource pairs (ctenophores feeding on copepods, interaction 12, and copepods consuming phytoplankton thin layers, interaction 4) fell just inside the lower-left quadrant, suggesting they should be unable to locate and exploit their respective resources. Only one consumer (the water flea *Bosmina* feeding on phytoplankton) fell in the upper-left quadrant, indicating a primary reliance on explosive reproduction to maintain populations within patches. Finally, most of the large marine predators exploited patches that are too ephemeral to allow reproduction within them.

As shown in Frost-Lessard space (Figure 4b), roughly half the consumers (18 out of 33, or 54%) had Le > 1 and were thus theoretically capable of depleting their resource patches. Of these patch-depleting consumers, all but two also had Fr > 1, suggesting they were able to effectively move between patches (Figure 4b). There was a positive correlation between Fr and Le in log-log space (r = 0.44, p = 0.01), with relatively faster-moving consumers also tending to consume their resource patches faster. Only two consumer-resource pairs (copepods-phytoplankton thin layers and ctenophores-copepods) fell in the upper-left quadrant of Figure 4b, where depletion of patches is possible but movement between them is not.

Across all consumer-resource pairs, there was also a positive correlation between the Str and Le numbers (r = 0.48, p = 0.004), indicating that the capacity for rapid population growth within a patch was associated with the ability to totally consume the resource (Figure 4c). However, consumer-resource pairs were present in all quadrants, so this tendency was far from a rule. Thirteen consumers fell in the upper-left quadrant (Str < 1, Le > 1), indicating they could deplete patches without being able to reproduce within them. Only three consumers--*Bosmina*, fruit flies, and elk--were capable of reproducing in a patch but incapable of depleting it, placing them in the lower-right quadrant with Str < 1 and Le < 1. Consumer populations will only persist if they can locate patches easily, reproduce within patches, or both, leading Grünbaum to predict that consumers must have either Fr or Str > 1. In our analysis, this prediction was borne out for 31 out of 33 interactions. The two exceptions were both planktonic consumers feeding on a planktonic resource: copepods - phytoplankton thin layers (Fr_dir_ = 0.2, Str = 0.35), and ctenophores - copepods (Fr_dir_ = 0.76, Str = 0.65). According to the logic of the dimensionless ratios, populations of these consumers should not be able to persist in nature--though this prediction is belied by the fact that ctenophores and copepods are two of the most widespread and abundant animal groups in the ocean. There are several possible explanations for this discrepancy. One is mismeasurement of the relevant patch scales or generation times: since both Frost and Strathmann numbers were within an order of magnitude of 1, only relatively small errors would be required to move them from above to below the critical value. Another, perhaps more likely, explanation is our use of generation times, rather than intrinsic rates of population growth, to estimate the Strathmann number. Both copepods and ctenophores can produce more than one offspring per generation, so their potential for explosive growth in patches is higher than predicted by the generation time alone. The intrinsic rate of increase would be a more appropriate basis for calculating the Strathmann number and should be used in future studies. However, values for it were not available in the literature for many of the consumers considered here, so we used generation time instead. The only consumer-resource interaction that fell in the upper-left quadrant in Fr-Str space was *Bosmina* - diatoms, another planktonic pair. Whether or not a purely reproductive strategy for patch exploitation is viable in non-planktonic interactions would be an interesting topic for future research.

This dimensionless framework reveals the importance of understanding a consumer’s movement ecology. Some consumers need directive search behavior to survive. Eight consumers would not be able to access their resource patches using diffusive search: white sharks, killer whales, chimney swifts, trout, common terns, common murres, green crabs, and the marine bacterium *Pseudoalteromonas*. It is striking that all but one of these (the green crab) feed on mobile prey in aquatic or aerial pelagic environments. This suggests that effective movement strategies are critical to the survival of these animals, and that this may be general to pelagic predators, whether they are pursuing prey through air or water. While we did not estimate costs of movement for consumers, the fact that none of these predators’ Frost numbers were >> 1 is consistent with Grünbaum’s conjecture that species will evolve movement abilities “good enough’’ to exploit their prey, but not greater.

### Dynamic similarities across scales and systems

A number of dynamic similarities were revealed when interactions were clustered according to their dimensionless ratios. We identified five groups of dynamically similar consumer-resource interactions (Figure 5). These groups should not be over-interpreted, since the selection of consumer-resource pairs and the number of clusters were both chosen semi-subjectively. The clusters could thus potentially shift with the addition of new pairs. Still, these clusters provide a useful perspective on generalities in patch dynamics, and hint at some of the reasons for their similarities and differences.

**Figure 5.**
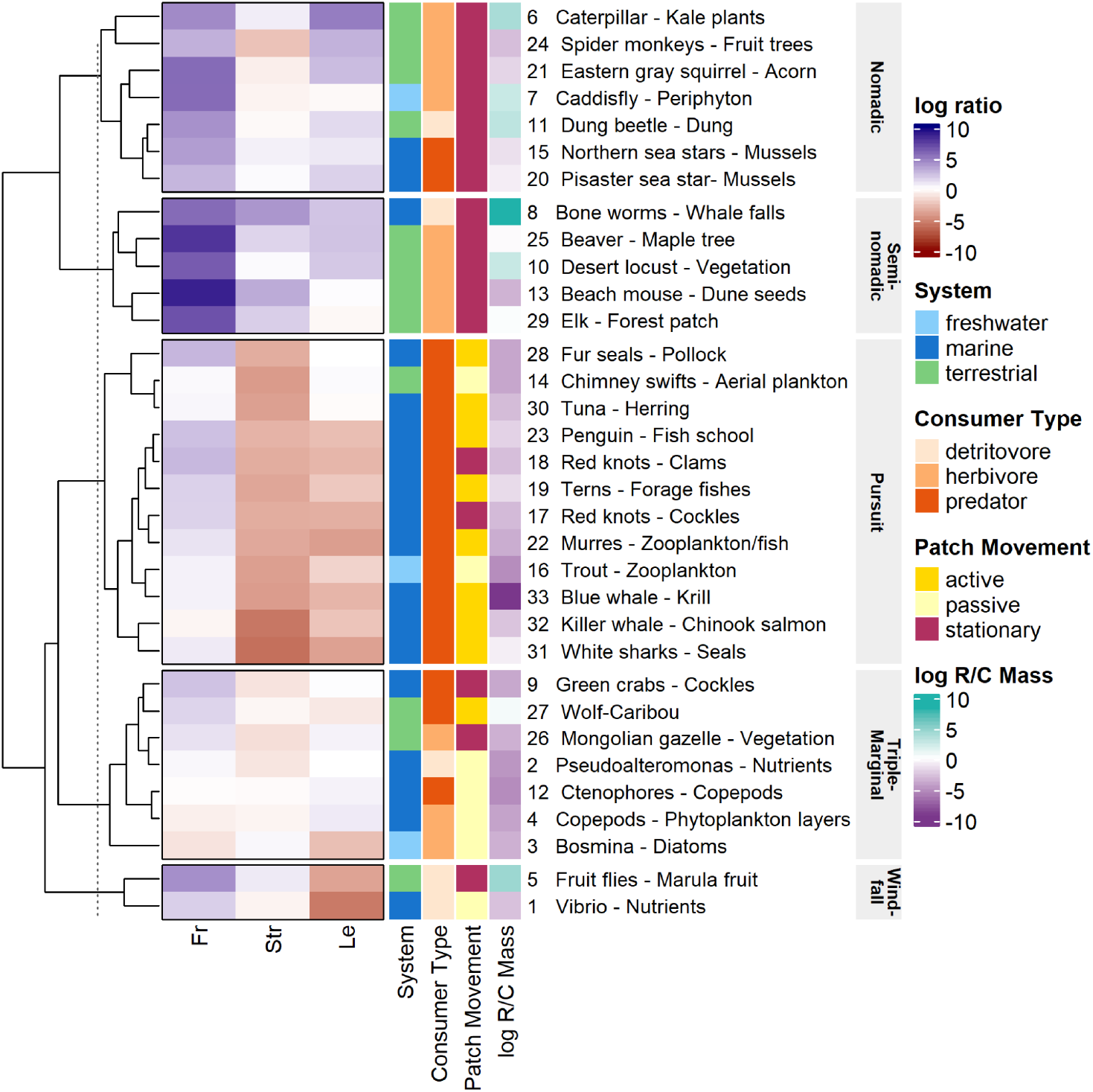
Clustering based on the log-transformed directed Frost, Strathmann, and Lessard numbers identified five groups of dynamically similar consumer-resource interactions. The three left columns in the heatmap (red-blue color scale) show the logarithmic values of Fr, Str, and Le for each interaction. The four right columns display the ecosystem in which the interaction takes place, the consumer and patch types, and the log-ratio of body mass between resource (R) and consumer (C). These latter four variables are shown for additional context but were not included in the clustering, which was driven entirely by Fr, Str, and Le. Each cluster is labeled with a descriptive name; see text for details.

The first cluster was defined by Fr > 1, Str ∼ 1, and Le > 1, indicating consumers could move relatively easily between patches and sometimes reproduce within them, but usually depleted them first. We termed these interactions *nomadic*, since these dynamics require constant movement by the consumers. We termed the second cluster *semi-nomadic*, since it shared similar features with nomadic interactions, though with faster movement and reproduction relative to patch duration (Fr >> 1, Str > 1) and a slightly lower chance of depletion (Le ∼ 1).

Both of these clusters were grouped together in the same branch of the dendrogram, and their broadly similar patch dynamics can be seen as falling on a continuum, with stationary resources exploited by mostly-herbivorous consumers. Consumers within the semi-nomadic cluster had, on average, higher Frost and Strathmann numbers, indicating they could locate and reproduce within patches more easily than the nomads. The nomadic cluster included three benthic aquatic and four terrestrial interactions, while all interactions in the semi-nomadic cluster were terrestrial except for *Osedax* bone worms on whale falls. Taken together, these two clusters encompass interactions where consumers travel easily between patches, moving from one to the next as they deplete their stationary resources. This depletion could happen relatively quickly (e.g., a quarter of an hour for a troupe of spider monkeys to eat the fruit off a tree) or over multiple generations (e.g., the decade or so for a population of bone worms to decompose a whale skeleton). The main difference between nomadic and semi-nomadic clusters was thus the length of residency the consumers had within their stationary resource patches.

The third cluster included 12 interactions. All were driven by movement (Fr > 1), with very slow reproductive dynamics (Str << 1) and marginal to very weak depletion (Str ∼ 1 to Str << 1). Based on their movement-dominated dynamics, we termed these interactions *pursuit-type*. While these consumers still moved relatively rapidly between resource patches, their Frost numbers were lower than those of the nomadic interactions (3 < Fr < 500). Pursuit-type consumers thus spent relatively more time searching for patches, perhaps because most of their resource patches moved actively, in contrast to the stationary patches in the nomadic and semi-nomadic interactions. The pursuit-type Strathmann and Lessard numbers were also uniformly < 1, meaning that patches were relatively shorter-lived and usually disappeared on their own before being fully exploited. Whereas the nomadic and semi-nomadic consumers could theoretically reproduce within patches, and would be forced to move only after depleting their resources, the pursuit-type consumers could never reproduce within patches. Instead, they relocated due to the patches’ inherent ephemerality.

Pursuit-type interactions exclusively involved carnivorous species preying on smaller-bodied animals. With the exception of chimney swifts – aeroplankton, all interactions took place in aquatic environments, and, except for red knots feeding on clams and cockles, all prey were mobile and suspended in a fluid medium. The separation of the pursuit-type and (semi-) nomadic groups highlights the dynamical differences between consumer-resource interactions in fluid environments and those in benthic or terrestrial environments where prey are attached to surfaces (Strathmann 1990; Steele 1991; Carr *et al*. 2003). For instance, the marine tuna - herring and terrestrial chimney swift - aeroplankton interactions were more similar to each other than either was to sea stars (a non-pelagic marine predator) or wolves (a non-aerial terrestrial predator). It is ironic that tuna, because of their voracity and pack hunting, have often been called the “wolves of the sea”—when in fact, they are better compared to an insectivorous bird, and wolves pursuing caribou are closer (at least in terms of patch dynamics) to crabs feeding on cockles (Figure 5).

The fourth cluster included 7 interactions, representing a varied collection of consumers, resources, and environments: microscopic and macroscopic; marine and terrestrial; carnivorous, herbivorous, and detritivorous. Their commonality was having all three dimensionless numbers close to the critical value of one—i.e., the timescales at which consumers moved between, reproduced within, and depleted patches were all similar to the patches’ durations. Consequently, these interactions were unlikely to satisfy any version of the mean-field assumption. Because movement, reproduction, and consumption processes were all marginal in these interactions, we called them *triple-marginal*.

For these consumers, small differences in reproductive or movement efficiency would mean the difference between effective and unsuccessful exploitation of patches. For instance, assuming diffusive movement, five of the consumers in this group would not be able to locate their resource patches, and based on the numbers we found, two consumers would be unable to locate patches even with directed movement. Several of the triple-marginal interactions *did* have large differences between the directed and diffusive Frost numbers, so if these consumers moved in a random-walk rather than directed fashion (as assumed for clustering), a spatial mean-field assumption might be justifiable. The triple-marginal cluster included four of the five planktonic consumers in our dataset (*Bosmina*, Copepods, Ctenophores, and *Pseudoalteromonas*), suggesting a hypothesis that planktonic consumers across a range of sizes may face similar challenges in gaining and maintaining access to resource patches. However, this cluster also included several non-planktonic interactions, including benthic and terrestrial predator-prey pairs and one terrestrial grazer.

The fifth cluster included only two interactions (fruit flies - marula trees and *Vibrio* - nutrients). Both had Fr > 1, Str ∼ 1, and Le << 1, indicating that the consumers could move between patches and possibly reproduce within them, but would not contribute significantly to resource depletion. Because these interactions involve consumers exploiting resources which appear suddenly and are more abundant than they can exploit, and because the fruit flies are literally consuming fallen fruit, we named them *windfall* interactions. These interactions were similar to the pursuit interactions in that movement between patches was possible and depletion was not (Fr > 1, Le << 1). However, the windfall interactions had Str ∼ 1, indicating that one generation of offspring could be produced within a nutrient plume or a fall of marula fruit before it diffused away or rotted, respectively. In these cases resource patches were both rich and moderately ephemeral, with a duration on the same order of magnitude as its consumer’s generation time. The small size of this cluster compared to the others is notable. While our selection of interactions cannot be assumed to be representative, the relative frequency of different types of consumer-resource interactions would be an interesting direction for future research.

### Are bacteria really gazelles?

Conceptualizing patchy consumer-resource dynamics in terms of dimensionless ratios can unite disparate ecological interactions into a common framework, and identify hidden similarities between them. However, these comparisons can at first appear quite abstract. To help make this style of thinking more concrete, we present a case study on one of the most surprising dynamic similarities we found, and the one that gives this paper its title: the similarity between Interaction 2, bacteria exploiting dissolved nutrients, and Interaction 26, gazelles grazing on the Mongolian steppe (Table 3, Figure 6).

**Figure 6.**
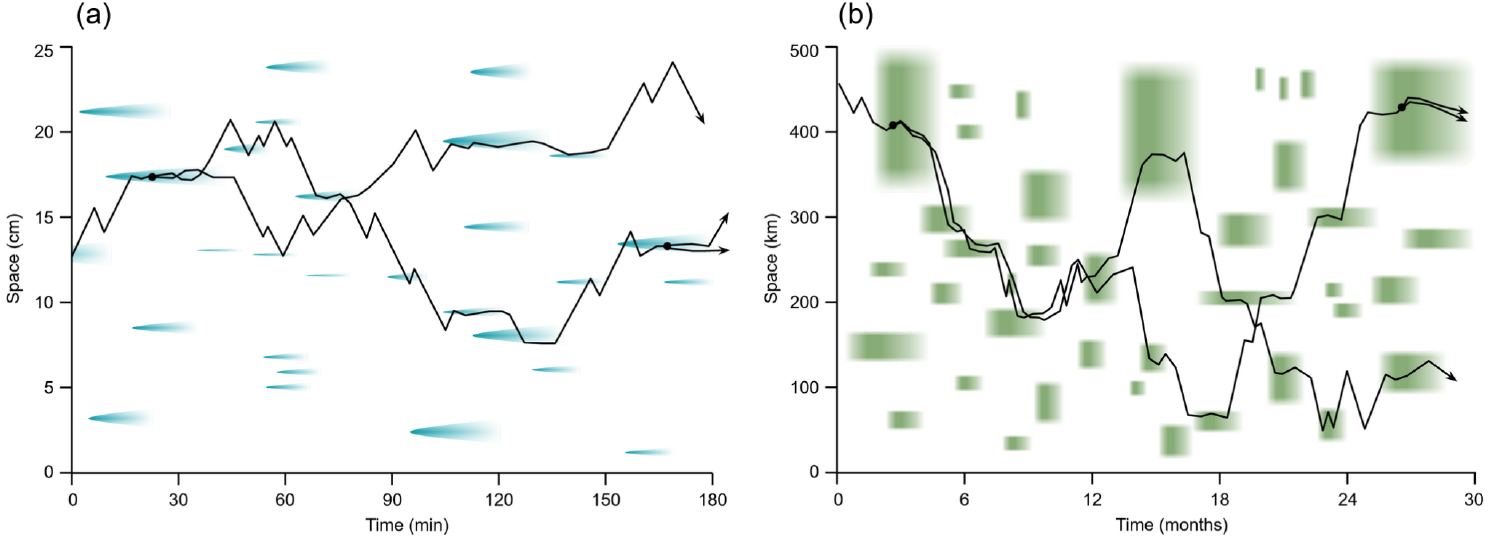
Patch exploitation by the marine bacterium *Pseudoalteromonas haloplanktis* and the Mongolian gazelle *Procapra gutturosa* are dynamically similar, despite differences in temporal scale on the order of 10^3^ and differences in spatial scale on the order of 10^6^. (a) Conceptual space-time diagram showing several bacteria (black lines) foraging for nutrient plumes from sinking phytoplankton cells (blue patches) in a simplified one-dimensional seascape. Black dots mark cell divisions, and hence the birth of new bacteria. (b) Similar diagram showing hypothetical paths of several gazelles browsing on patches of steppe vegetation which bloom following localized rains. As in a), black dots mark births of new organisms. The spatial and temporal scales of the patches are approximately accurate for each system, as are the speeds and generation times of the organisms. Note that for simplicity only a few trajectories are plotted; in reality the density of both organisms within their patches would be significantly higher.

*Pseudoalteromonas haloplanktis* is a cold-water planktonic bacterium which, in this interaction, seeks out a plume of nutrients left behind by a dead, sinking phytoplankton cell. Such plumes are typically separated by several cm, and disperse via diffusion over about 15 minutes. Laboratory tracking has shown that *P. haloplanktis* can swim incredibly fast for its body size, moving up nutrient gradients at speeds of up to 0.4 mm/s (Barbara & Mitchell 2003), implying a directed Frost number of 2.0. Their doubling time of ∼3 hours (Piette *et al*. 2010) is fast by human standards, but 12 times longer than the duration of an individual nutrient plume (implying Str = 0.083), meaning that a bacterium dividing in a plume cannot expect its daughter cells to inherit the resource patch. Still, the relatively high density of bacteria that can aggregate in such a nutrient plume means that they can absorb the nutrients at about the same rate as diffusion carries the nutrients away, leading to a Lessard number approximately equal to one.

The Mongolian gazelle, *Procapra gutturosa*, differs in several respects from *P. haloplanktis*: to name a few, it is terrestrial, reproduces sexually, and possesses a body made of multiple eukaryotic cells that is 2.8 quadrillion times the mass of *P. haloplanktis*. Nevertheless, the foraging dynamics of gazelle are quantitatively similar, with (directed) Fr = 12.3, Str = 0.04, and Le = 3.0, all on the same order of magnitude as corresponding numbers for the bacterium. An examination of the gazelles’ natural history provides an explanation for this similarity: Herds of gazelles travel nearly constantly across the steppe, tracking patches of productive pasture that appear following rains. These patches last on the order of 30 days, long enough for herds to locate them, but an order of magnitude shorter than the gazelles’ 2-year generation time. Still, the size and density of the herds (281 gazelles km^-2^, Olson *et al*. 2009) mean that they actively graze down their pastures.

The bacterium and the gazelle’s interactions were closer to each other in logarithmic Fr-Str-Le space than to any others considered here. Both consumers are fast for their size and highly mobile relative to their resources and would be expected to spend the majority of their time inside patches. While these patches are long-lasting enough to enable easy discovery, they disappear too quickly to host population growth, meaning that each consumer must visit 10-20 patches per generation. At typical consumer densities, each patch is a finite resource, being depleted faster than it would be in the absence of the consumers. Figure 6 shows the conceptual space-time arrangement of patches and consumer trajectories for both these interactions. The dynamic similarity between them is intuitively obvious when comparing the two panels: if one ignores the axis labels and stylized patch shapes, the distribution of patches and the consumers’ space-time trajectories are difficult to tell apart.

Of course, this is a deliberately simplified picture of these interactions’ dynamics. In many respects, bacteria are *not* gazelles. Nutrient plumes are small compared to the spaces separating them, while patches of steppe vegetation vary widely in size and connectivity (Mueller *et al*. 2008). The consumers’ sensory capabilities and foraging behaviors also differ significantly. While *Pseudoalteromonas* can perform chemotaxis, gazelles use multiple senses, memory, and social information to locate food. Bacterial division occurs whenever cellular development and nutrient assimilation permit, while gazelles have a seasonal reproductive cycle. Finally, the bacteria move in a three-dimensional environment, whereas the gazelles are restricted to the (approximately) flat surface of the earth. Nevertheless, dimensional analysis suggests that some of these differences in biology will be less important than dynamic similarities between the arrangement of patches in space and time, and the ways consumers travel between and exploit them over their lives’ courses. Outside this dimensionless scaling framework, such a surprising dynamic similarity would not be apparent.

### Significance, perspective, and future directions

As is the case for most dimensionless ratios, Fr, Str, and Le are based on a simplified caricature of the real world, and do not capture all dynamical similarities (or differences) that can exist between systems (e.g., density dependence, functional responses, other species interactions, responses to environmental drivers; Rogers & Munch 2020). Further, each set of numbers characterizes an *interaction*, not a taxon. The same consumer could have different values for Fr, Str, and Le when feeding on a different resource, or even the same resource in a different place or time. Just as the nondimensional numbers can identify dynamic similarities across species, they can also identify dynamic differences within a single species as it grows, changes behavior, or encounters different resource and environmental conditions. In reality, resources are heterogeneous at a range of spatiotemporal scales, and patches can be nested inside each other. It is thus important to think critically about which of these scales are relevant for the consumers and the ecological questions at hand when calculating these dimensionless ratios. Examining the sensitivity of Fr, Str, and Le to patch scales in hierarchical resource landscapes would be a worthwhile direction for future investigation. Ultimately, though, it is important to remember that these ratios are not intended to be more precise than an order of magnitude: they are tools for reasoning about the relative magnitudes of different rates, not precisely modelling dynamics.

As a general framework, we believe dimensionless ratios are widely useful. Although the ratios explored here were derived from specific details about individual consumers and resources, clear groupings emerged that reflected similarities in life history (herbivorous vs. predatory consumers, mobile vs. stationary resources) and environment (“pelagic” vs. “benthic” systems). These separations hint at fundamental tradeoffs long discussed in ecology and evolution (Hutchinson 1961; Menge & Sutherland 1987; Strathmann 1990). The dimensionless ratio approach is similar to the use of functional traits within community ecology (e.g., McGill *et al*. 2006), where the diversity of organism traits are reduced to their functional similarities in a comparative framework. Dimensionless descriptions highlight how species interact with each other and their environment and provide an opportunity for ecologists studying dynamically similar systems to learn from each other. It may also suggest experimentally tractable systems that can be used as proxies for dynamically similar interactions that occur at experimentally intractable spatial and temporal scales.

While our selection of consumer-resource pairs was extensive, it was not random or representative. For instance, our author group has little expertise with insects, so they are underrepresented in this paper. Additionally, we did not consider plants or fungi as consumers, even though they move between generations via seed or spore dispersal and can actively seek resource patches by extending roots or mycorrhizae. Finally, we found it surprisingly difficult to locate values for basic rates (speeds, reproductive and consumption rates, patch sizes, densities, etc.) in the literature, even for well-studied species, and most values we did find were published in older papers. While funders, publishers, and researchers may not consider measuring and reporting basic natural history information as “high-impact” or career-advancing ecology, their value to future researchers is hard to overstate (Greene 2005).

Understanding when patchiness matters and when it may be ignored is a constant challenge to ecological modellers. The Frost, Strathmann, and Lessard numbers can serve as diagnostic tools to assist in model development by identifying when mean-field approaches will work for a given system. For theoreticians seeking to understand patchy consumer-resource dynamics (e.g., Hein & Martin 2019), empirical information on the Fr-Str-Le space occupied by real organisms can aid in model formulation and placement of realistic parameter bounds. The broad utility of these values is analogous to the Reynolds number in fluid dynamics: it is not the *only* number one needs to design an airplane, or predict the weather, but neither is possible without it. To model and understand ecological patchiness, we ought to start from a common quantification. The Frost, Strathmann, and Lessard numbers, proposed by Grünbaum (2012) and quantified here, may provide such a starting point from which to develop a deeper, more general understanding of patch dynamics in ecology.

## Acknowledgements

This paper grew out of discussions at the Ecological Dissertations in the Aquatic Sciences symposium (Eco-DAS) at the University of Hawaii at Manoa in October 2018 and was supported by funding from the National Science Foundation (OCE-1356192) and the Association for the Sciences of Limnology and Oceanography. We are grateful to Dr. Paul Kemp and Kristina Remple for their work organizing the symposium, and to all our Eco-DAS 2018 friends for the stimulating discussions and good times. MLS was supported by a grant from the Simons Foundation (Award ID 602015). Gemma Carroll, Stephen Katz, Peter Dudley, and Patrick Ressler provided early feedback which greatly improved the manuscript. Finally, we would like to thank Daniel Grünbaum for supplying the ideas and framework that inspired this effort.

## Author contributions

SSU conceived the study. All authors discussed the study’s design, reviewed literature, and curated data. ANC, SSU, and TLR analyzed the data. ANC and SSU drafted the manuscript with input from all authors. ANC, CCS, JSS, SSU, and TLR contributed to figures and tables. All authors revised and edited the manuscript.

## Supplementary material

**Figure S1.**
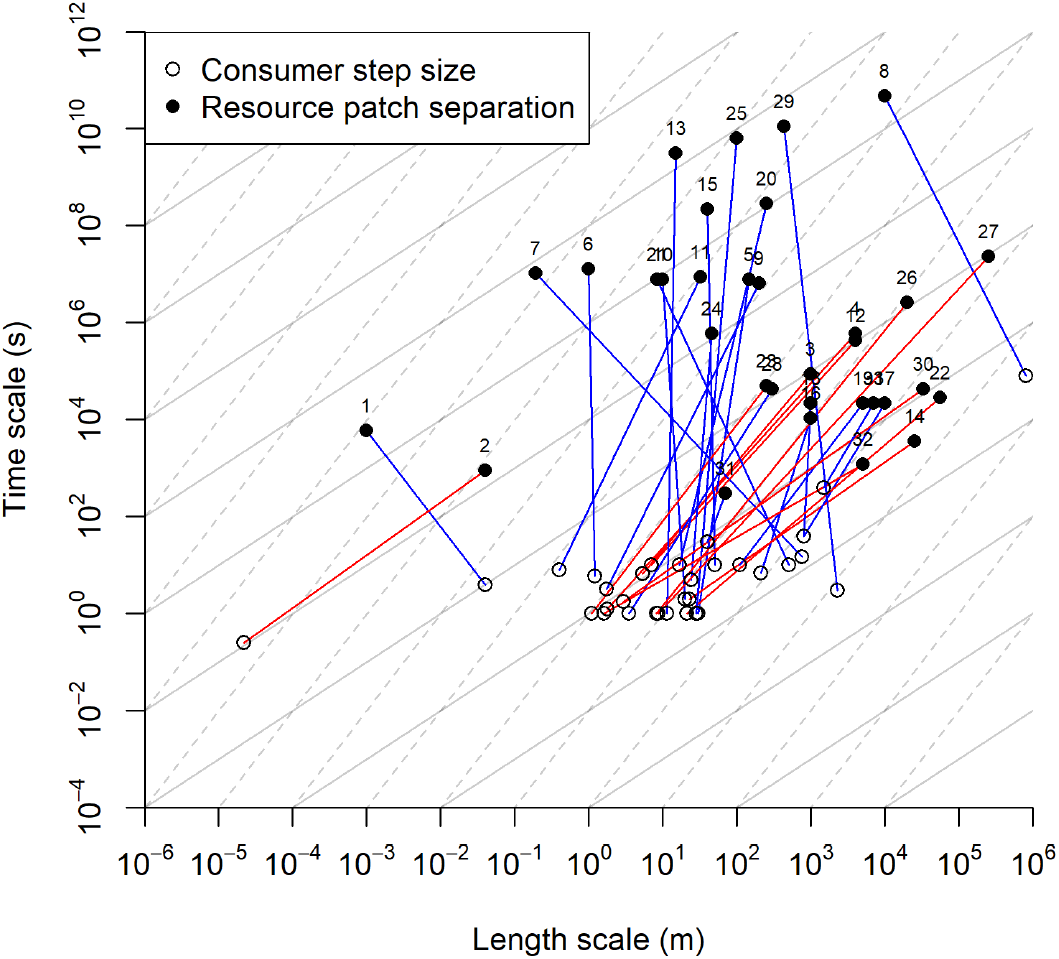
Patch availability by diffusive versus active movement. Critical Frost numbers for diffusive and directed searches, based on Grünbaum’s (2012) Figure 1a. Filled points show the duration and separation scales of a resource patch. Each of these patch points is connected to an open point showing the “minimum step” of its consumer, placed at the turning interval τ and the distance traveled between turns at its representative velocity *s* (Table 3). Numbered labels correspond to the consumer-resource interactions listed in Table 3 and all figures in the main text. A consumer can only find patches if its Fr ≳ 1. On logarithmic length-time axes, the critical Frost numbers for directed and diffusive motion correspond to lines with slopes of 1 or 2, respectively. The grid of solid and dotted lines in the background shows these slopes for reference. If the slope of the segment connecting a consumer to its resource is less than 1, the resource is unavailable. If the slope is between 1 and 2 (red lines), patches can be located by a directed search, but not a random search. If the slope is > 2 (blue lines), patches are available with any search strategy.

**Figure S2.**
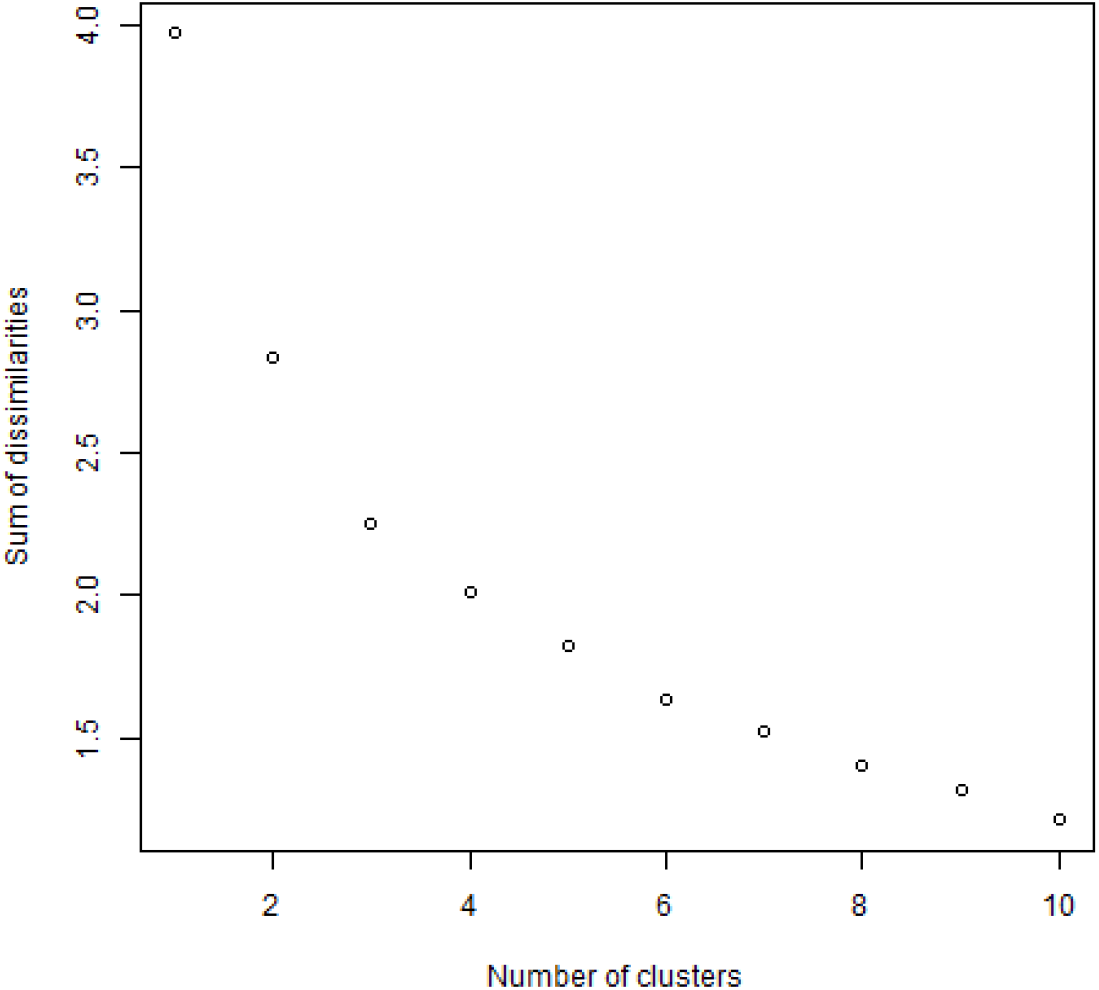
Elbow plot used to select the number of clusters.

**Table S1**. Complete presentation of rates and dimensionless ratios for all consumer-resource pairs. Parameters are defined in Table 2. An abbreviated version of these data is given in Table 3. Consumer and resource quantities are presented in their original units from the primary literature, which vary from interaction to interaction. For instance, the “currency” of an interaction could be expressed in individuals or biomass, densities could be areal or volumetric, etc. However, when plugged into the equations in Table 2, the quantities within each row are dimensionally consistent. Each quantity is listed with its source in the literature, as both a URL permalink/DOU and author-year citation (cf. references below). Where the value of the quantity was not simply reported in the source paper, requiring some interpretation or calculations on our part, these are summarized in the “_comment” column.

